# Genomic relatedness strengthens genetic connectedness across management units

**DOI:** 10.1101/130138

**Authors:** Haipeng Yu, Matthew L. Spangler, Ronald M. Lewis, Gota Morota

## Abstract

Genetic connectedness refers to a measure of genetic relatedness across management units (e.g., herds and flocks). With the presence of high genetic connectedness in management units, best linear unbiased prediction (BLUP) is known to provide reliable comparisons between genetic values. Genetic connectedness has been studied for pedigree-based BLUP; however, relatively little attention has been paid to using genomic information to measure connectedness. In this study, we assessed genome-based connectedness across management units by applying prediction error variance of difference (PEVD), coefficient of determination (CD), and prediction error correlation (r) to a combination of computer simulation and real data (mice and cattle). We found that genomic information (**G**) increased the estimate of connectedness among individuals from different management units compared to that based on pedigree (**A**). A disconnected design benefited the most. In both datasets, PEVD and CD statistics inferred increased connectedness across units when using **G**- rather than **A**-based relatedness suggesting stronger connectedness. With r once using allele frequencies equal to one-half or scaling **G** to values between 0 and 2, which is intrinsic to **A**, connectedness also increased with genomic information. However, PEVD occasionally increased, and r decreased when obtained using the alternative form of **G**, instead suggesting less connectedness. Such inconsistencies were not found with CD. We contend that genomic relatedness strengthens measures of genetic connectedness across units and has the potential to aid genomic evaluation of livestock species.

The problem of connectedness or disconnectedness is particularly important in genetic evaluation of managed populations such as domesticated livestock. When selecting among animals from different management units (e.g., herds and flocks), caution is needed; choosing one animal over others across management units may be associated with greater uncertainty than selection within management units. Such uncertainty is reduced if individuals from different management units are genetically linked or connected. In such a case, best linear unbiased prediction (BLUP) offers meaningful comparison of the breeding values across management units for genetic evaluation (e.g., Kuehn et al., 2007).

Structures of breeding programs have a direct influence on levels of connectedness. Wide use of artificial insemination (AI) programs generally increases genetic connectedness across management units. For example, dairy cattle populations are considered highly connected due to dissemination of genetic material from a small number of highly selected sires. The situation may be different for species with less use of AI and more use of natural service mating such as for beef cattle or sheep populations. Under these scenarios, the magnitude of connectedness across management units is reduced and genetic links are largely confined within management units.

Pedigree-based genetic connectedness has been evaluated and applied in practice (e.g., Kuehn et al., 2009; Eikje and Lewis, 2015). However, there is a relative paucity of use of genomic information such as single nucletide polymorphisms (SNPs) to ascertain connectedness. It still remains elusive in what scenarios genomics can strengthen connectedness and how much gain can be expected relative to use of pedigree information alone. Connectedness statistics have been used to optimize selective genotyping and phenotyping in simulated livestock (Pszczola et al., 2012) and plant populations (Maenhout et al., 2010), and in real maize (Rincent et al., 2012; Isidro et al., 2015), and real rice data (Isidro et al., 2015). These studies concluded that the greater the connectedness between the reference and validation populations, the greater the predictive performance. However, 1) connectedness among different management units and 2) differences in connectedness measures between pedigree and genomic relatedness were not explored in those studies. For better understanding of genome-based connectedness, it is critical to examine how the presence of management units comes into play. For instance, genomic relatedness provides relationships between distant individuals that appear disconnected according to the pedigree information. In addition, it captures Mendelian sampling that is not present in pedigree relationships (Hill and Weir, 2011). Thus, genomic information is expected to strengthen measures of connectedness, which in turn refines comparisons of genetic values across different management units. The objective of this study was to assess measures of genetic connectedness across management units with use of genomic information. We leveraged the combination of real data and computer simulation to compare gains in measures of connectedness when moving from pedigree to genomic relationships. First, we studied a heterogenous mice dataset stratified by cage. Then we investigated approaches to measure connectedness using real cattle data coupled with simulated management units to have greater control over the degree of confounding between fixed management groups and genetic relationships.

## Materials and Methods

### Mice data

We analyzed a heterogeneous stock mouse population established for quantitative trait mapping (Valdar et al., 2006; Solberg et al., 2006). It was originally derived from eight inbred strains (DBA/2J, C3H/HeJ, AKR/J, A/J, BALB/cJ, CBA/J, C57BL/6J, and LP/J), followed by 50 generations of pseudorandom mating. This process introduced recombinants that allow high-resolution mapping (Valdar et al., 2006; Solberg et al., 2006). This population was used for one of the first empirical applications of genomic selection in animals (Legarra et al., 2008) and later used for an array of quantitative genetic studies. The data consisted of 1,884 individuals from 169 full-sib families with approximately 11 siblings per family. Each individual was genotyped with 10,946 SNPs yet none of the full-sib parents were genotyped. We removed SNPs with a minor allele frequency (MAF) less than 0.05, resulting in 10,339 markers for analysis. The mice were reared in 523 cages or management units that created shared environments. The majority of full-sibs were housed in the same cages and distributed to three cages on average, i.e., a full-sib family was typically reared together in three cages. Pedigree relationships within and across full-sib individuals were 0.5 and 0, respectively. This resulted in an extreme case of genetic disconnectedness across management units. Thus, the extent of connectedness was determined by the presence or absence of full-sibs in different management units.

### Cattle data

Pedigree information of dairy cattle was available on 1,929 cattle collected over six generations starting from a base generation 0 to generation 5 (Wimmer et al., 2015). Among those, 500 individuals, mostly coming from generations 2 and 3 (> 90%), had both phenotypes and genotypes. Historic pedigree information in addition to the 500 individuals are a source of connectedness as the pedigree-based relationship matrix was constructed from the entire pedigree. The 500 individuals were genotyped for 7,250 SNP markers. The average missing rate of genotypes across the entire SNPs was 0.0002. We imputed missing genotypes by sampling alleles from a Bernoulli distribution with the marginal allele frequency used as a parameter. We retained 6,714 SNPs after removing markers with MAF less than 0.05. We simulated management units in two steps: 1) individuals were clustered and 2) clusters were assigned to management units. The k-medoid clustering was performed to cluster individuals into distinctive groups. In particular, we used partitioning around medoids, which is considered a robust version of K-means (Kaufman and Rousseeuw, 1990; Reynolds et al., 2006). We formed sets of clusters so that individuals in the same groups were more similar to each other than to those in other groups. We selected the number of clusters by optimum average silhouette width algorithm implemented in the cluster and fpc R packages. This algorithm minimizes dissimilarity measures among individuals within the same cluster using the Euclidean metric and finds the optimal number of clusters that returns the lowest average dissimilarity computed from each cluster. The clustering was based on the **A** matrix, which was converted to a dissimilarity matrix by calculating the distance from the highest similarity to each similarity value in such a way that the relationship with the largest value becomes zero. We simulated the four following scenarios.

- Scenario 1: Completely disconnected - all clusters allocated to their own management units
- Scenario 2: Disconnected - one-half of clusters allocated to management unit 1 and remaining half assigned to management unit 2
- Scenario 3: Partially connected - approximately one-third of clusters allocated to management unit 1, another one-third to management unit 2, and the remaining one-third of clusters assigned to both managements to act as a link to connect the two managements units indirectly
- Scenario 4: Connected - all clusters equally allocated to the two management units

Subsequently, appropriate incidence matrices were constructed and we computed connectedness statistics across management units employing pedigree and genomic relationships.

### Prediction error variance

Genetic connectedness statistics are typically defined as a function of the inverse of the coefficient matrix. For instance, Kennedy and Trus (1993) proposed a genetic connectedness measure as the average prediction error variance (PEV) of the difference in predicted genetic values between all pairs of individuals in different management units. The PEV can be obtained from Henderson’s mixed model equations (MME) (Henderson, 1984). We constructed MME according to a standard linear mixed model **y = Xb + Zu + ϵ**, where **y** is a vector of phenotypes, **X** is an incidence matrix of management units, **b** is a vector of effects of management units, **Z** is an incidence matrix relating individuals to phenotypic records, **u** is a vector of random additive genetic effects, and **ϵ** is a vector of residuals. The phenotypic vector **y** was standardized to have mean of 0 and variance of 1 so that results can be compared across different scenarios. The variance-covariance structure for this model is

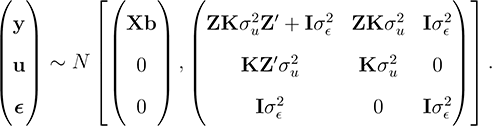

where 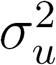 is the genetic variance, 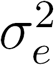 is the residual variance, and **K** is a positive (semi)definite relationship matrix defined later.

The inverse of the MME coefficient matrix is represented as

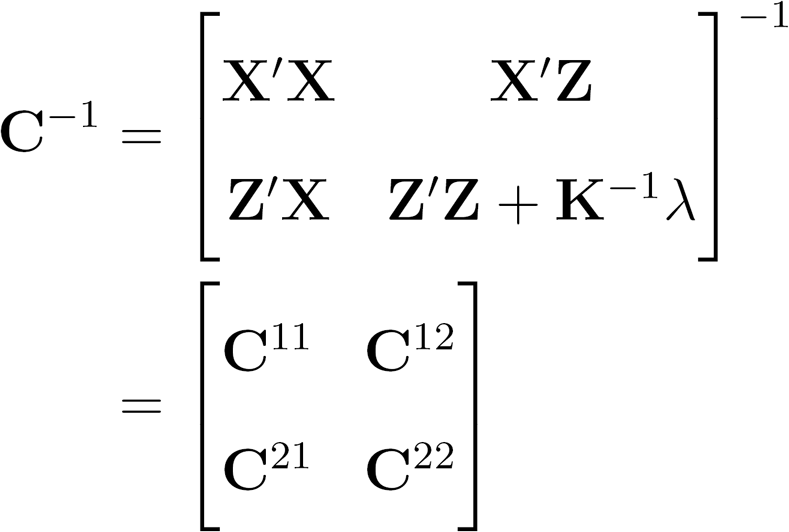

where λ is the ratio of variance components 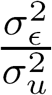. The PEV of genetic value for the *i*th individual (*û_i_*) is given by

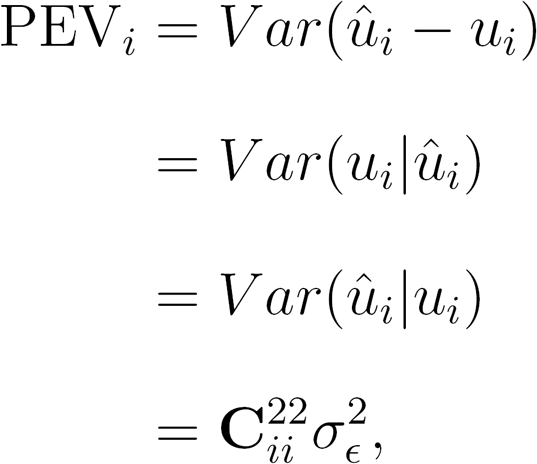

where 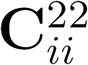 is the *i*th diagonal element of C^22^ coefficient matrix. Note that PEV can be interpreted as the proportion of additive genetic variance not accounted for by the prediction. Equivalently, the matrix of PEV can be computed as

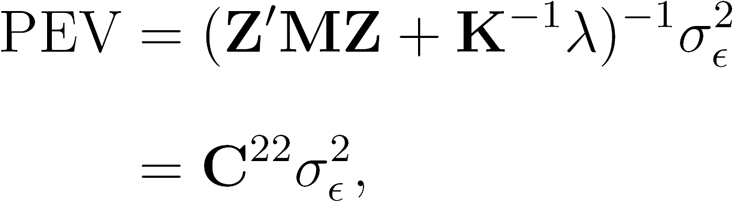

where **M** is the absorption (projection) matrix for fixed effects where **M = I – X(X′X)^-^X′**, which is orthogonal to the vector space defined by X (i.e., MX = 0). This avoids calculating the inverse of the entire coefficient matrix, which is useful when the number of columns of X is large or analysis involves repeated computation of PEV.

### Genetic connectedness

We computed three genetic connectedness statistics: the PEV of the difference (PEVD) between genetic values (Kennedy and Trus, 1993), the coefficient of determination (CD) of the difference between predicted genetic values (Laloë, 1993), and the prediction error correlation (r) between genetic values of individuals from different management units (Lewis et al., 1999). The first two statistics were originally used to evaluate the accuracy of individual estimated breeding values (EBV) and later extended to assess inherent risk in comparing individuals across management units. First, genetic connectedness between two individuals, *i* and *j*, was measured as PEVD (Kennedy and Trus, 1993)

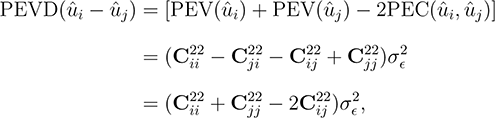

where PEC_*ij*_ is the prediction error covariance or covariance between errors of genetic values, which is off-diagonal element of the PEV matrix. If PEVD is small, individuals are said to be connected. The idea behind using PEVD as a measure of connectedness is that the accurately estimated genetic values of individuals have smaller PEV and that the pairs of genetically related individuals in the different management units have a positive prediction error covariance. Throughout this study, we used a scaled PEVD following Kuehn et al. (2008) by scaling PEVD by the additive genetic variance to express connectedness without units of measurement.

Similarly, CD is closely related to PEVD and is defined by scaling the inverse of the coefficient matrix by corresponding coefficients from the relationship matrix. We can view CD as the squared correlation or reliability between the predicted and the true difference in the breeding values (Laloë et al., 1996). This is given by

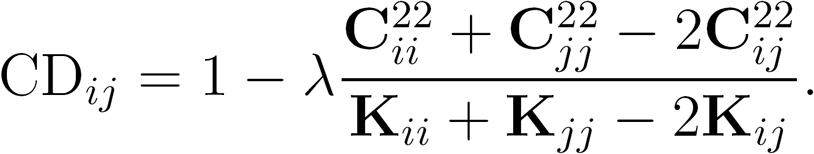

or pairwise comparison. In contrast to PEVD, CD accounts for the reduction of connectedness due to relationship variability between individuals under comparison. This statistic is bounded between 0 to 1, with larger values indicating increased connectedness.

The r is obtained by transforming a PEV matrix into predictive error correlation matrix. For individuals *i* and *j*, this statistic is given by

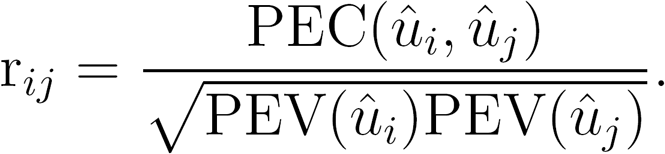

The rationale behind r is that there is no connectedness when PEC is zero (Lewis et al., 1999). Similar to CD, r is also bounded between 0 and 1. The larger the r, the greater the connectedness.

### Connectedness summary

We can generalize connectedness between any pair of management units *i*′ and *j*′ by setting up a corresponding contrast vector x that sums to zero (i.e., **1′x** = 0) (Laloë, 1993). The PEVD of contrast x in genetic values is given by

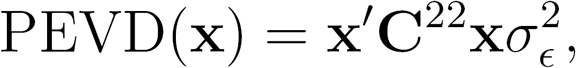

where x is a column vector including *1/n_i′_*, *−1/n_j′_*, and 0, for the elements corresponding to *i′*th unit, *j′*th unit, and the remaining units, respectively, where *n_i′_* and *n_j′_* were the numbers of individuals belonging to *i*′th and *j*′th units, respectively. In a contrast vector notation, pairwise CD between management units *i′* and *j*′ is given by

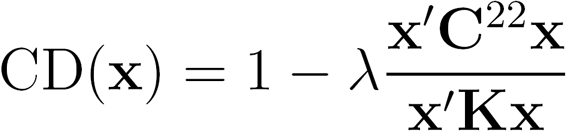

For the r statistic, a similar summary statistic can be derived as

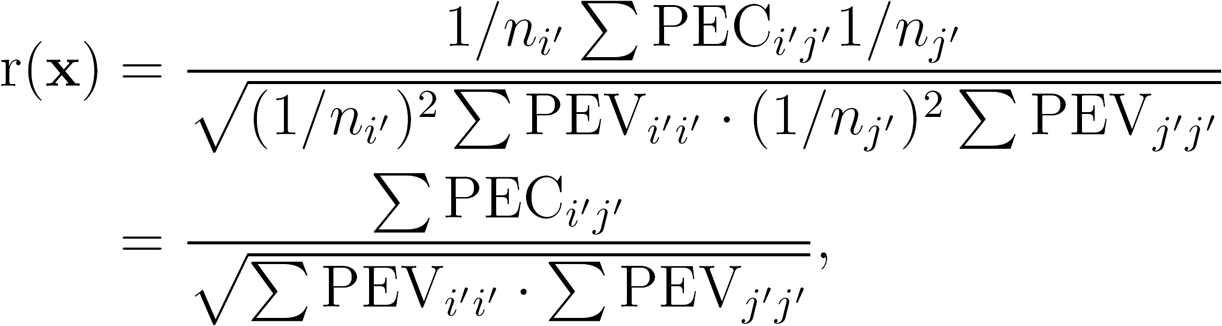

where 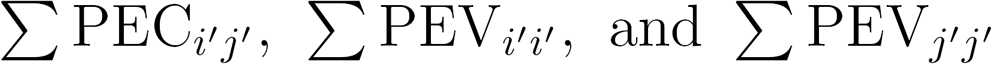 were the sums of the elements of PEC*_i′j′_*, PEC*_i′i′_*, and PEC*_j′j′_* respectively (Kuehn et al., 2008). However, in the Appendix we show that when this summary statistic is applied across units it provides a reasonable summary for a pedigree relationship matrix but it is not suitable for a genomic relationship matrix when the total number of management units is two. Thus, we reported connectedness by averaging the r statistic for all pairs of individuals in across management units.

### Relationship matrix

Connectedness is realized through a genetic relationship matrix under the BLUP framework. Three genetic connectedness statistics defined above require information about covariance structures among individuals or genetic values that evaluate relatedness. We considered five *n* × *n* relationship kernel matrices (**K**) in this study, where n is the number of individuals. The numerator relationship matrix, **K** = **A**, is based on relatedness due to expected additive genetic inheritance. This can be computed directly from pedigree information, and reflects the probability that alleles are inherited from a common ancestor and thereby are identical by descent (IBD). The off-diagonal elements are twice the kinship coefficients and are equivalent to the numerators of Wright’s correlation coefficients (Wright, 1921, 1922). The majority of genetic connectedness literature is based on the pedigree relationship matrix, i.e., average relationships assuming conceptually, an infinite number of loci. On the other hand, the genomic relationship matrix, **K** = **G**, captures genomic similarity among individuals. The matrix **G** is a function of the matrix of allelic counts (*ω_i,j_ ∈* 0,1, 2), where *i* = 1, …, *n* and *j* = 1, …, *m* denote the indices of individuals and of markers, respectively. Each element of the allele content matrix **W** is the number of copies of the reference allele. Under Hardy-Weinberg equilibrium, 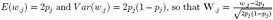 is a standardized incidence matrix of allelic counts, where *p_j_* is the allele frequency at the *j*th marker. The **G** matrix is constructed from a crossproduct of scaled marker genotype matrix **W** divided by some constant, i.e., the number of markers under assumption of unity marker variance

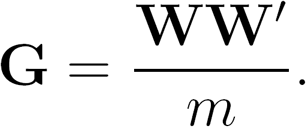

The standardization of **W** and the constant in the denominator make the **G** matrix analogous to the **A** matrix (VanRaden, 2008). This genomic relationship matrix estimates the proportion of the genomes of two individuals that is identical by state (IBS).

One concern that arises when comparing the **A** and **G** matrices is that these two matrices are not on the same scale. The G matrix represents the estimate of a covariance (correlation) structure among individuals marked by SNPs with the potential having some negative off-diagonal entries. Such negative values indicate that some individuals are molecularly less related than average pairs of individuals in the sense of IBS if the population were in Hardy-Weinberg equilibrium (e.g., Toro et al., 2002). This mostly happens when the current population is defined as a base population, namely, computing the **G** matrix by using the estimates of observed allele frequencies from the current population (Powell et al., 2010). While the negative coefficients arising from IBS can be interpreted as negative correlations of alleles (Toro et al., 2002), this is contrast to the **A** matrix which is defined as an IBD. In the **A** matrix, a founder population is assumed to be the unselected base population. This may impact some of the connectedness statistics used in this study. For this reason, we also considered two other genomic relationship matrices: a **G_0.5_** matrix and a scaled **G** matrix, **G_s_**, so that the genomic relationship matrix is on nearly the same scale as the A matrix. The **G_0.5_** matrix was created by scaling the W by 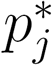, instead of *p_j_*, where 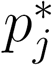 is the estimate of allele frequency in the base population. Because allele frequencies in the base population are unknown, we set all 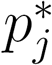 equal to 0.5 under the assumption of no selection (VanRaden, 2007; Toro et al., 2011; Vitezica et al., 2011). The **G_0.5_** matrix constructed in this way does not create any negative coefficients for the both mice and cattle datasets. The correlations between **G** and **G_0.5_** (defined as correlation between elements of upper triangular matrix including diagonals) were 0.81 and 0.98 for mice and cattle, respectively.

Alternatively, a min-max scaler, one of the common scaling methods, was employed to scale the **G** matrix. The min-max scaler transforms inputs into the given range of minimum and maximum values. The scaled genomic relationship between *i*th and *j*th individual was given by

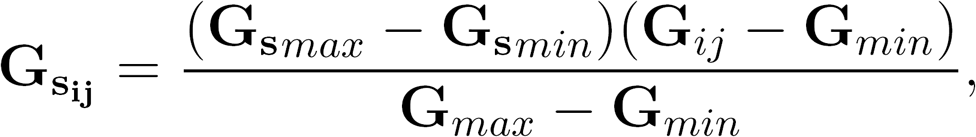

where **G***_min_* and **G***_max_* are the minimum and maximum elements of **G**, and **G***_ij_* is the *i*th, *j*th element of **G**. The **G**_**s***min*_ and **G**_**s***max*_ define the range of minimum and maximum values of elements of **G_s_**. These values were set to 0 and 2, respectively, according to the minimum and maximum values of numerators of Wright’s correlation coefficients. This scaling sets negative off-diagonal entries in the G matrix to 0 (Momen et al., 2017). Note that the correlation between G and G_s_ is equal to one because a correlation is invariant to changes in scale.

Lastly, the covariance between ungenotyped and genotyped individuals was jointly modeled through a hyrbrid matrix where **K** = **H**. The H matrix can be viewed as a matrix that combines pedigree and genomic relationships. By considering the distribution of genetic values of ungenotyped individuals conditioned on genetic values of genotyped individuals, it can be shown (Legarra et al., 2009; Christensen and Lund, 2010) that

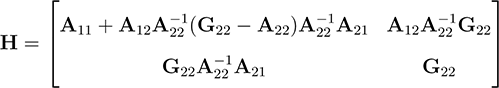

where **A**_11_, **A**_12_(**A**_21_), and **A**_22_ are numerator relationship matrices among ungenotyped, ungenotyped and genotyped, and genotyped individuals, respectively. **G**_22_ = **G, G_0.5_**, or **G_s_** is the genomic relationship matrix for genotyped individuals. In addition to **A, G, G_0.5_**, and **G_s_**, the **H** matrix was used for the cattle dataset that spans several generations. We treated individuals at generations three, four, and five as genotyped individuals and earlier generations as ungenotyped individuals. This reflects a practical situation in typical breeding programs, where the majority of genotyped individuals are concentrated in more recent generations. This partitioning resulted in 65% ungenotyped and 35% genotyped individuals, simulating a realistic scenario where there are more ungenotyped than genotyped individuals (e.g., Legarra et al., 2009).

### Principal component analysis of measures of connectedness

Principal component analysis (PCA) of PEVD, CD, and r pairwise individual-based matrices computed under the four different simulated scenarios in the cattle dataset was used to cluster individuals. The prcomp function in R was used to produce principal component scores and the principal component (PC) plots were generated with the ggbiplot package based on the first two PC.

### Heritability

For simulation, we used two heritability values (*h*^2^ = 0.8 and *h*^2^ = 0.2) by varying the ratio of variance components 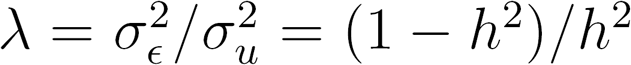 assuming an animal model, where 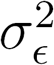 and 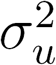 are residual and genetic variances, respectively.

### Data availability

The mouse dataset is available at http://wp.cs.ucl.ac.uk/outbredmice/heterogeneous-stock-mice/ and the cattle dataset is downloadable from the synbreedData R package at https://cran.r-project.org/web/packages/synbreedData/index.html.

## Results

### Mice data

#### Absence of full-sibs

The average (standard deviation) of pedigree relationships among individuals in the same management units was 0.491 (0.058) because of the aforementioned full-sib family assignments. The genomic counterpart (**G**) gave a similar estimate of 0.494 with a slightly increased standard deviation of 0.087 due to Mendelian sampling variation (Hill and Weir, 2011). The average across management unit pedigree-based genetic connectedness was 1.299 when measured by PEVD and *h*^2^ = 0.8 (Table 1). Measures of connectedness increased using genomic data (**G**) by reducing PEVD to 0.456. With *h*^2^ = 0.2, while the overall genetic connectedness decreased, genomic information (**G**) lowered PEVD compared to that of pedigree. Use of the **G_0.5_** reduced PEVD more than that of the G, hence increased the measures of connectedness. Using the scaled genomic relationship matrix increased connectedness statistics compared to those of the pedigree-based, but they were less than those with **G**. Similarly, use of **G** matrix compared to the **A** matrix strengthened measured connectedness in CD for both *h*^2^ = 0.8 and *h*^2^ = 0.2. The **G_0.5_** matrix also increased measures of connectedness compared to those of the **A** and the **G_s_** matrix resulted in the greatest measures of connectedness among the four relatedness matrices. Both PEVD and CD statistics confirmed that genome-wide markers increased the degree of connectedness measured between individuals within the across management units. However, the connectedness measures assessed by r were less when the **G** was compared with the **A**. On the other hand, the **G_0.5_** and the scaled genomic relationship matrix **G_s_** showed greater connectedness measures than those of the **A**.

**Table 1:**
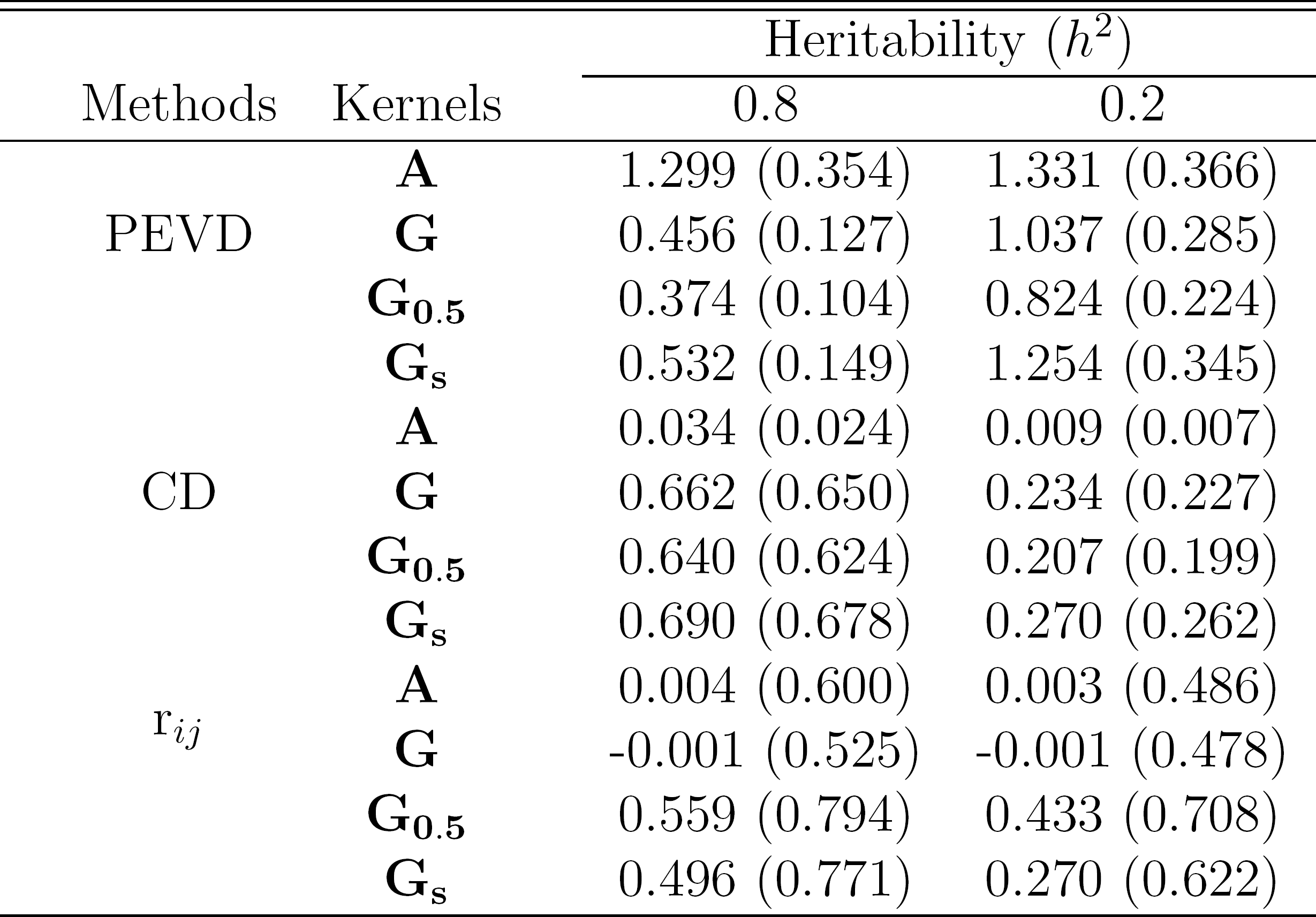
Average genetic connectedness measures across management units in the mice data. PEVD, CD, and r denote prediction error variance of the difference, coefficient of determination, and prediction error correlation. We compared pedigree-based **A**, standard genome-based **G**, genome-based **G_0.5_** assuming equal allele frequencies, and scaled genome-based **G_s_** matrices to evaluate relationships among individuals. Two heritability values 0.8 and 0.2 were simulated. Values inside parentheses represent connectedness when at least one full-sib pair was present in different management units.

| Methods | Kernels | Heritability ($h^2$) | |
| --- | --- | --- | --- |
|  |  | 0.8 | 0.2 |
| PEVD | $\mathbf{A}$ | 1.299 (0.354) | 1.331 (0.366) |
| | $\mathbf{G}$ | 0.456 (0.127) | 1.037 (0.285) |
| | $\mathbf{G}_{0.5}$ | 0.374 (0.104) | 0.824 (0.224) |
| | $\mathbf{G}_s$ | 0.532 (0.149) | 1.254 (0.345) |
| CD | $\mathbf{A}$ | 0.034 (0.024) | 0.009 (0.007) |
| | $\mathbf{G}$ | 0.662 (0.650) | 0.234 (0.227) |
| | $\mathbf{G}_{0.5}$ | 0.640 (0.624) | 0.207 (0.199) |
| | $\mathbf{G}_s$ | 0.690 (0.678) | 0.270 (0.262) |
| $r_{ij}$ | $\mathbf{A}$ | 0.004 (0.600) | 0.003 (0.486) |
| | $\mathbf{G}$ | -0.001 (0.525) | -0.001 (0.478) |
| | $\mathbf{G}_{0.5}$ | 0.559 (0.794) | 0.433 (0.708) |
| | $\mathbf{G}_s$ | 0.496 (0.771) | 0.270 (0.622) |

#### Presence of full-sibs

The increased estimates of disconnectedness was less when at least one full-sib was present in different management units for PEVD. For instance, comparisons between absence or presence of full-sibs across management units were 1.299 vs. 0.354 and 0.456 vs. 0.127 for pedigree-based vs. genome-based (**G**) PEVD, respectively. The presence of full-sibs in different management units decreased PEVD. However, corresponding statistics for CD were lower with the existence of full-sibs. This is explained by the fact that CD penalizes the estimates of connectedness when genetic variability is small. The CD statistic attempts to decrease the average PEV of the contrast while maintaining the variability of relatedness. Laloë (1993) stated that increased estimate of connectedness should not be achieved by simply using genetically similar individuals and CD is the most relevant connectedness statistic in terms of genetic progress of agricultural species. This was confirmed in the mice data illustrating that the presence of full-sibs decreased the estimates of CD. Regardless of absence or presence full-sibs across-units, genomic information elucidated additional relationships, thus increasing connectedness relative to pedigree. This trend was also true for the **G_0.5_** and **G_s_** matrices. With r, when transitioning from **A** to **G**, the values of the statistic reduced; however, the **G_0.5_** and **G_s_** yielded greater values of connectedness than those of pedigree in the existence of full-sibs. In all cases, using one of the **G, G_0.5_**, or **G_s_** matrix increased the estimates of connectedness statistics as compared to using the **A**. As shown in Table 1, we found a similar overall pattern when *h*^2^ was set to 0.2, although connectedness remained less than the alternative higher heritability. Replacing pedigree with genome-wide markers increased the degree of connectedness captured among individuals in disconnected management units.

#### Illustrative examples

To illustrate how **G** matrix impacted our measures of connectedness, we chose five management units including full-sib and non-full-sib individuals. In this example, management units ”19F”, ”29A”, and ”36F” share at least one pair of full-sib individuals, whereas management units ”12A” and ”13C” do not share any full-sib individuals across management units. Figure 1 shows PEVD-derived connectedness across management units when *h*^2^ = 0.8. Comparison across management units with full-sibs in common had smaller PEVD hence greater connectedness. The molecular information captures more of the genetic connectedness relative to pedigree for across management units. We further investigate how the **G** or **G_s_** increased connectedness across management units using PEVD and r by examining the specific components in the scaled PEV matrix by following similar examples in Kennedy and Trus (1993). For instance, disconnected ”19F” and ”13C” management units contained five full-sib individuals each. The following matrix contains the pedigree-based PEV for the 10 individuals.

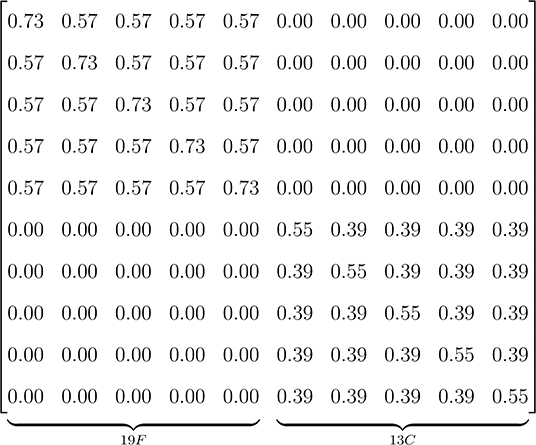

The first five individuals belong to ”19F” and the remaining individuals belong to ”13C”. Because there are no full-sib pairs across management units, off-diagonals are all zero. Thus, PEVD and r of two individuals across management units are 0.73 + 0.55 – (2 × 0) = 1.28 and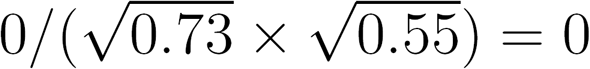, respectively.

When the **A** matrix is replaced with the **G** matrix, the PEV matrix becomes

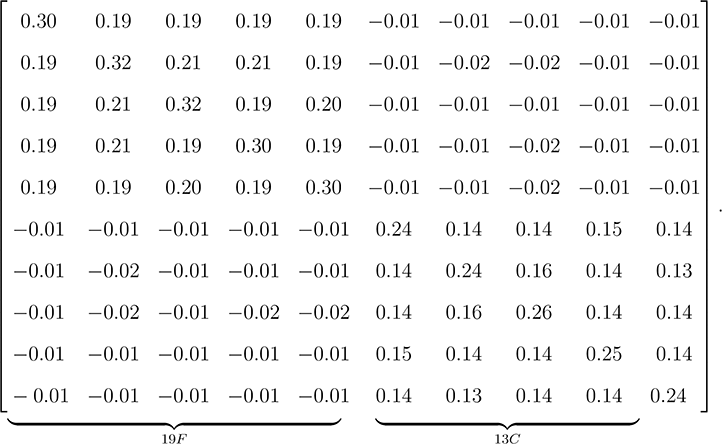

Average genomic relationships within management units were 0.419 and 0.440 for individuals in “19F” and “13C”, respectively, whereas across management unit genomic relationships were -0.09. The off-diagonals of zeros in pedigree-based PEV were replaced with small negative values. Although the diagonal elements within management units are not all equal because of Mendelian sampling, PEVD between the first individuals from respective management units are 0.30 + 0.24 – (2 × −0.01) = 0.56. Given that off-diagonal elements are negligible, the rate of PEVD reduction from shifting from **A** to **G** is almost 50% with the most of difference coming from decreased PEV in the diagonals. Specifically, the rates of PEV reduction (diagonals) from **A** to **G** were 59% and 56% for the first individuals in “19F” and “13C”, respectively. The rates of PEC reduction cannot be defined since all of the off-diagonal elements are zeros in **A**. Note that the r statistic using the **G** matrix does not yield increased estimates of connectedness compared with that using **A** because 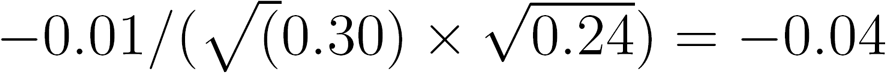. Now consider the **G_s_** matrix

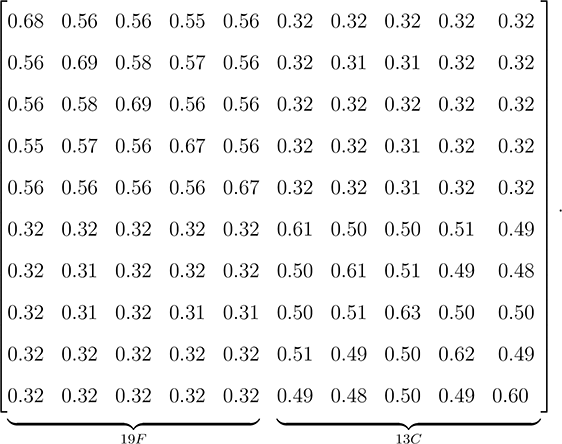

Here the PEV matrix within management units more closely resembles those from pedigree-based PEV. In addition, the negative elements across management units PEV were replaced with positive values. With use of **G_s_**, PEVD and r between the first individuals from the two management units are 0.68 + 0.61 — (2 × 0.32) = 0.65 and 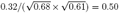, respectively. When the scaled genomic relationship matrix **G_s_** is used, r yields an increased connectedness estimate as compared to using pedigree-based relationships.

**Figure 1:**
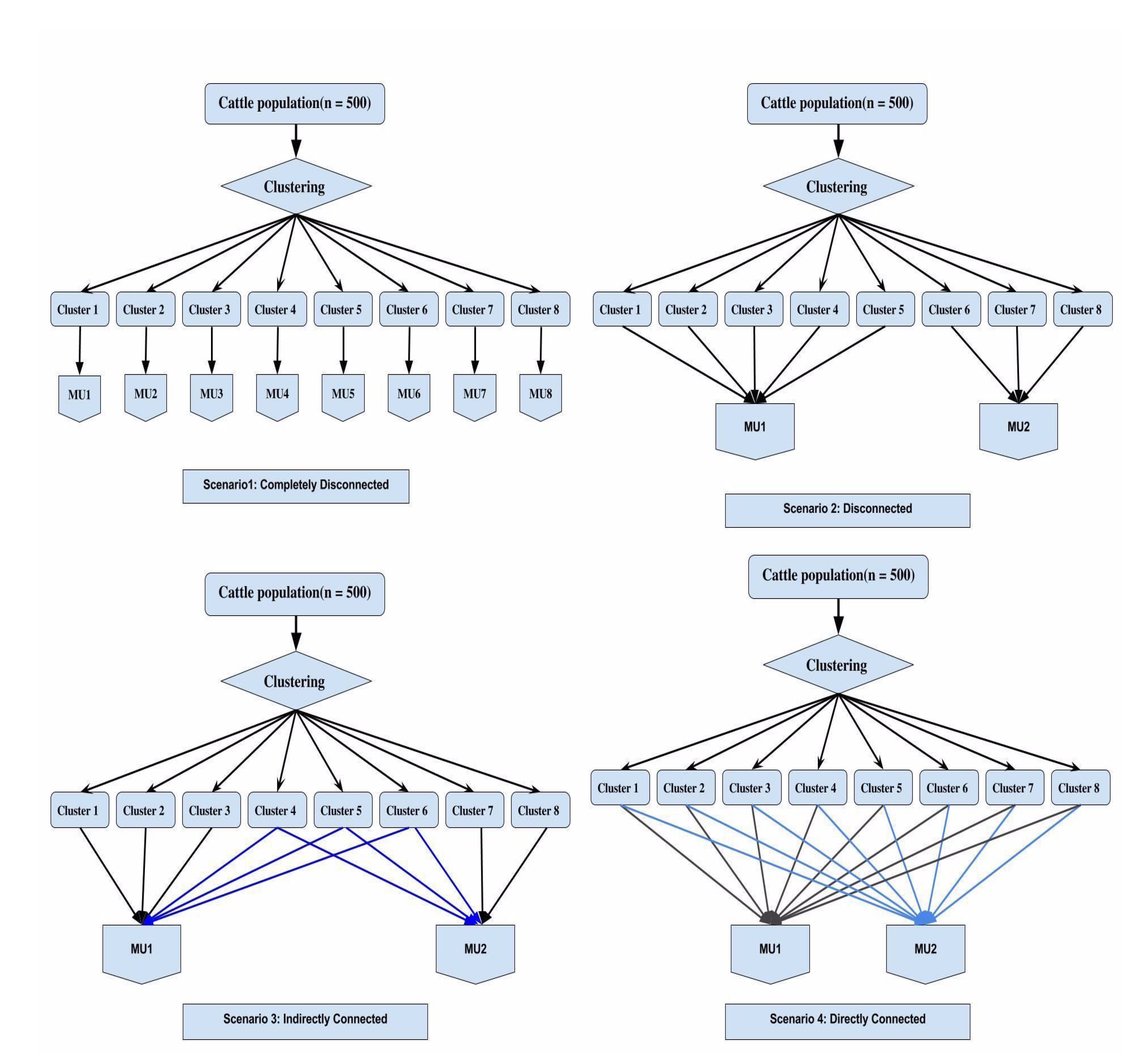
Prediction error variance of the difference (PEVD) for across five management units in the mice dataset. Management units ”19F”, ”29A”, and ”36F” share at least one pair of full-sibs individudals with each other, whereas ”12A” and ”13C” do not share any individuals across management units. The left and right are pedigree-based (**A**) and genomic-based (**G**) connectedness, respectively. Darker color represents less genetic connectedness.

Likewise, a subsequent example with “19F” and “36F” which can be viewed as connected management units (Figure 1). Each management unit contains five full-sibs as in the previous case. The following matrix of pedigree-based PEV includes these 10 individuals.

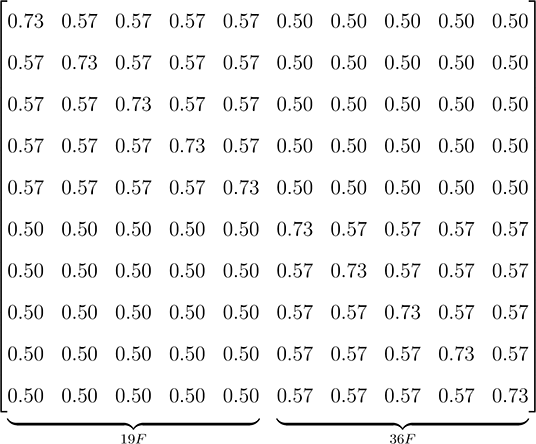

The first five individuals belong to “19F” and the remaining individuals belong to “36F”. In this case, off-diagonals are non-zero due to the presence of shared full-sib information across management units. Here PEVD and r of two individuals is 0.73 + 0.73 – (2 × 0.50) = 0.46 and 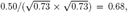, respectively. Relative to the pedigree-based non-full-sib comparison, a significant increase in connectedness was observed. The majority of the increase in connectedness is due to increased PEC between individuals.

The following is the PEV matrix when pedigree is substituted with the genome-wide markers from **G**.

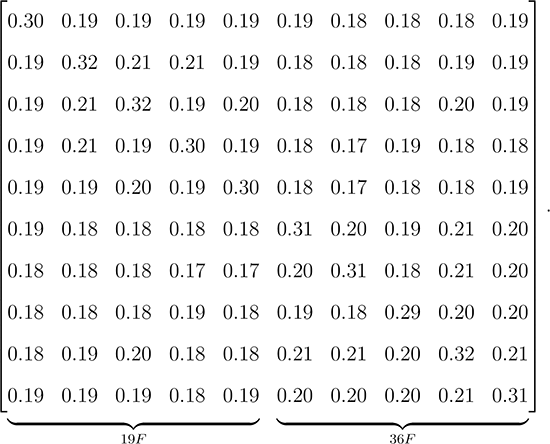

Average genomic relationship within “36F” was 0.465 whereas average genomic relationship across “19F” was 0.419. The off-diagonals no longer have small negative values and all elements of PEV were reduced in comparison to those using A. Here PEVD and r between the first individuals from the two management units are 0.30 + 0.31 – (2 × 0.19) = 0.23 and 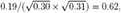, respectively. Again, while genomic information increased estimates of connectedness as measured by PEVD, that was not the case for r. The reduction in PEVD from A to G was about 50% and both diagonals and off-diagonals contributed to increasing connectedness estimates. In particular, the rates of PEV reduction (diagonals) from A to G were 59% and 58% for the first individuals in “19F” and “36F”, respectively. The reduction in PEC due to the use of G was 62% for these two individuals, which was larger than the reduction of the diagonals. This contributed to the unexpected results for r because this statistic is based on a ratio. Now consider use of the scaled genomic relationship matrix, **G_s_**, which yielded the following PEV matrix

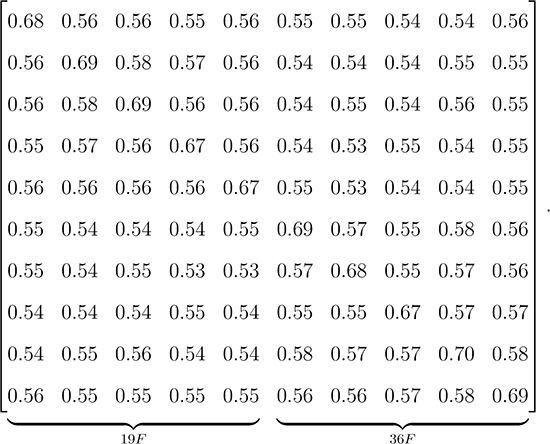

The PEV matrix including both within and across management units are more analogous to those of the pedigree-based PEV. Here PEVD and r between the first individuals from the two management units are 0.68 + 0.69 – (2 × 0.55) = 0.27 and 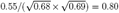, respectively. The result reaffirms that scaling **G** to be on the same scale as **A**, genomic relatedness leads to an increases in the ratio sensitive connectedness statistic, a change consistent with the other statistics. Collectively, these particular examples, suggest that genomic information provided by **G** or **G_s_** changes the estimates of relationship coefficients among animals and refines the estimates of connectedness.

### Cattle

#### Clustering

The partitioning around medoids clustering method yielded eight clusters. Table 2 contains descriptive statistics for those clusters. The number of individuals per cluster varied from 36 to 127. The average of within cluster pedigree-based relationships was around 0.05 except for cluster 6, in which distant relatives were grouped together. Between clusters, all pedigree-based relationships were close to zero. Each cluster was assigned to management units in four simulated scenarios as summarized in Figure 2.

- Scenario 1: Each cluster was assigned to its own management unit
- Scenario 2: Clusters 1, 2, 3, 4, and 5 were assigned to management unit 1 and clusters 6, 7, and 8 were assigned to management unit 2
- Scenario 3: Clusters 1, 2, 3 were assigned to management unit 1, clusters 7 and 8 were assigned to management unit 2, and individuals in clusters 4, 5, and 6 were assigned to both management units 1 and 2 to act as link among clusters or individuals that partially connect the two management units
- Scenario 4: Individuals in clusters 1 to 8 were equally assigned to management units 1 and 2

**Table 2:** Descriptive statistics of the 8 clusters in the cattle data.

| Cluster | Number of individuals | Average pedigree relationship |
| --- | --- | --- |
| 1 | 52 | 0.054 |
| 2 | 61 | 0.053 |
| 3 | 46 | 0.040 |
| 4 | 36 | 0.052 |
| 5 | 43 | 0.043 |
| 6 | 127 | 0.005 |
| 7 | 55 | 0.055 |
| 8 | 80 | 0.047 |

**Figure 2:**
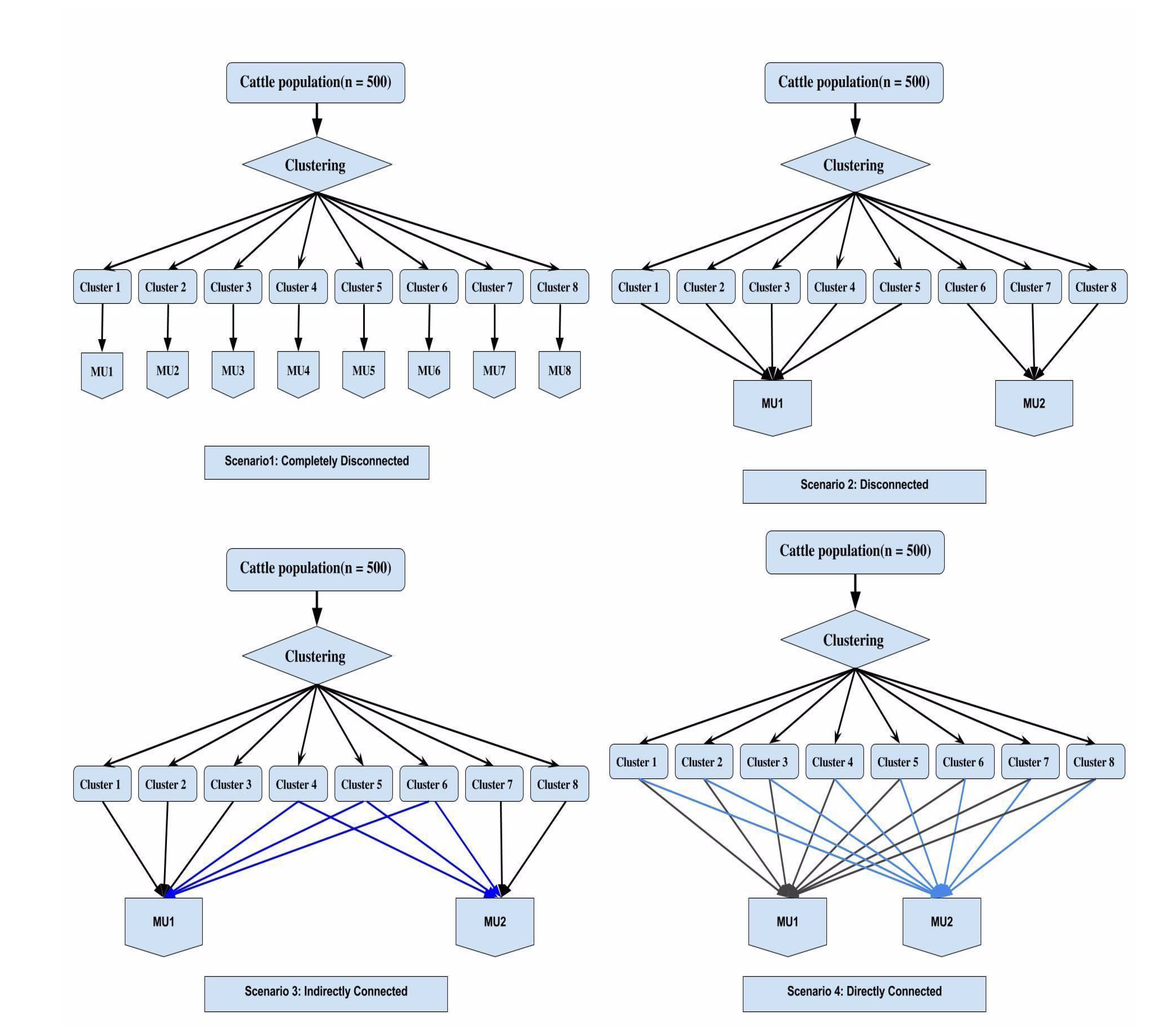
Four simulation scenarios considered in the cattle dataset. MU stands for management unit. Scenario 1: Completely disconnected -8 clusters assigned to separate management unit. Scenario 2: Disconnected - clusters 1, 2, 3, 4, and 5 assigned to management unit 1 and clusters 6, 7, and 8 assigned to management unit 2. Scenario 3: Partially connected - clusters 1, 2, 3 assigned to management unit 1, clusters 7 and 8 assigned to management unit 2, and the remaining clusters 4, 5, and 6 assigned to both management units 1 and 2 that act as link among clusters or individuals that partially connect the two management units. Scenario 4: Connected - all clusters 1 to 8 were equally assigned to management units 1 and 2.

The number of individuals in management units 1 and 2 were approximately equal in scenarios 2, 3, and 4. We computed PEVD, CD, and r for each of the four scenarios and compared genetic connectedness when using the **A, G, G_s_**, and **H** kernel matrices.

#### Prediction error variance of the difference

Across management unit PEVD for each of four scenarios are presented in Tables 3. Connectedness estimates increased across management units when transitioning from scenario 1 to scenario 4 for both heritability levels. Figure 3 shows the relative increase of genetic connectedness as measured with PEVD, as a percentage, across management units in comparison to scenario 1. Genetic connectedness across management units in scenario 1 was compared to across management unit connectedness obtained from scenarios 2, 3, and 4. We observed increased genetic connectedness as more individuals from the same clusters were shared between management units, resulting in the highest connectedness estimates in scenario 4. Transitioning from scenario 1 to scenario 4 increased connectedness for **A** and **G** for both heritability levels. The proportional increase in genetic connectedness in pedigree-based relationships were larger than those of genomic-based relationships because **G** matrix substantially increased measured connectedness between disconnected management units in scenario 1, reducing the gains in the following scenarios 2, 3 and 4. Also, as heritability increased, larger values of connectedness were observed. In general, with **G** and **G_0.5_** increased the estimates of connectedness compared to those of the **A** regardless of heritability levels. This is in agreement with the mice dataset. However, with **G_s_**, values of PEVD were unexpected: although with scaled **G_s_** produced estimates of connectedness that are higher than those with **A** when *h*^2^ was set to 0.8, the same pattern was not observed for *h*^2^ = 0.2.

**Figure 3:**
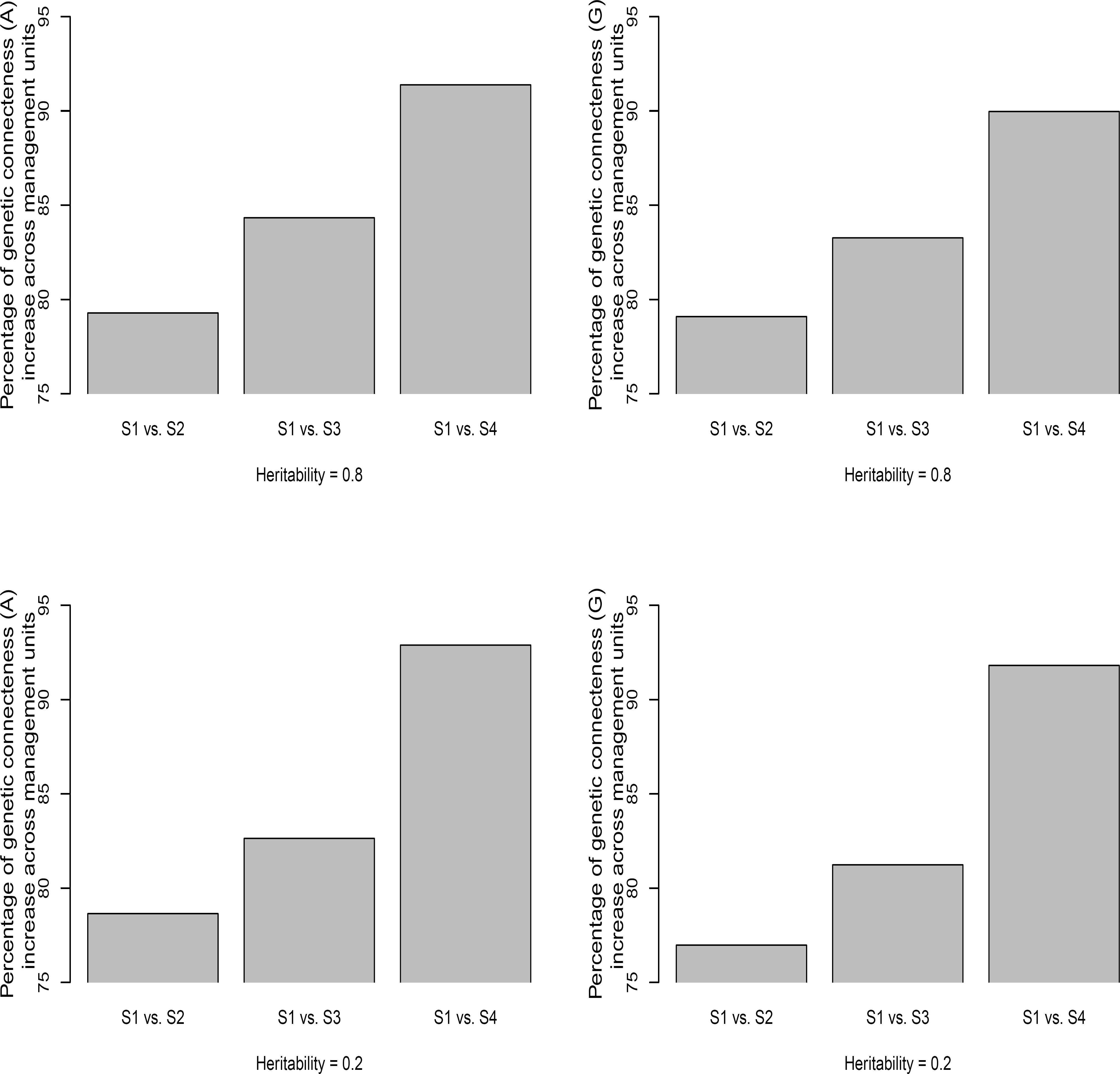
Percentage of relative increase in prediction error variance of the difference (PEVD) across management units in comparison to base Scenario 1. Two heritability values 0.8 and 0.2 were simulated. S1 (completely disconnected), S2 (disconnected), S3 (partially connected), and S4 (connected) represent four management unit scenarios. Left: A matrix. Right: G matrix.

#### Coefficient of determination

Across CD for each of four scenarios are presented in Tables 3. Similar to PEVD the extent of connectedness across management units increased when moving from scenario 1 to scenario 2 and 3 regardless of the heritability levels. Supplementary material Figure S1 in File S1 shows the percentage increase in CD across management units when scenario 1 was treated as a base comparison. As with PEVD, CD statistics revealed an increase in the degree of connectedness as more individuals from the same clusters were assigned to different management units. However, the increase of CD was not observed when transitioning from scenario 1 to scenario 4. Again, this is because CD accounts for the reduction of connectedness due to reduced relatedness variability between individuals under comparison in scenario 4. This pattern was observed for both pedigree and genomic-based connectedness. Overall, **G, G_0.5_**, and **G_s_** all produced CD greater than those with **A** regardless of heritability level, yielding consistent measures of connectedness.

#### Prediction error correlation

Prediction error correlations across management units for each of four scenarios are presented in Tables 3. The results align with those of the mice dataset in that **G**-based r statistics behave erratically in all scenarios making them difficult to interpret. However, the anticipated increases in r were observed with transition from the **A** to the **G_0.5_** or the scaled **G_s_** matrix. Here **G_0.5_** and **G_s_**-based measures consistently yielded greater connectedness values than those of pedigree counterparts. Figure S2 in File S1 shows the percentage increases in r across management units when scenario 1 was treated as a base comparison. Here the **G_s_** instead of the **G** matrix was used. The results align with those of PEVD and CD, where the extent of pedigree-based and genomic-based r statistics increase the most when more individuals from the same clusters were assigned to different management units. The magnitude of the increase was larger when heritability was greater. However, the increase of connectedness moving from Scenario 1 to 2 was not observed in pedigree-based measures. While both scenarios are not connected designs because pedigree-based relationships across the eight clusters were close to zero, it is interesting to note that with pedigree-based r scenario 2 was more disconnected than scenario 1. From Kennedy and Trus (1993), this is because stronger within unit connectedness can reduce between unit connectedness.

#### Ungenotyped and genotyped individuals

We considered a scenario where only individuals in younger generations were genotyped in the cattle dataset. For this purpose, we used the **H** matrix that blends ungenotyped and genotyped individuals. As shown in Table S1 in File S1, results using the H matrix lie somewhere between the results obtained when using the **A, G**, and **G_s_** matrices. This is expected because the **H** matrix was created from a combination of **A** and **G** or **G_s_**. Although an increase in measures of connectedness was observed compared to using the pedigree alone, this increase was smaller than when all individuals were genotyped. This finding suggests the possibility of strengthening the degree of connectedness even when only a subset of individuals was genotyped. An exception was observed when **H** constructed from **G_s_** for PEVD: in this case the measures of connectedness were less than that from **A**.

#### Principal component analysis of connectedness

Principal component plots for CD derived from **A** and **G** matrices for scenarios 1 and 4 are presented in Figures 4 and 5, respectively. These correspond to the two extreme scenarios considered in the cattle dataset. In scenario 1 with *h*^2^ = 0.8, eight clusters assigned to distinctive management units were separated from each other as expected using pedigree-based relationships (Figure 4). Genomic information brought these eight clusters closer to each other, thus shortening the distance between individuals from different management units. While eight clusters were less distinguishable from one another due to lower heritability, the same pattern was observed when *h*^2^ was 0.2. These findings align with the fact that use of genomic information increases measures of connectedness compared to pedigree. In both cases, cluster 6, which consisted of unrelated individuals, was clustered far away from the other clusters in the pedigree-based analysis. PCA yielded two clear clusters in scenario 4 when *h*^2^ = 0.8, which correspond to the two management units considered (Figure 5). Replacing **A** with **G** resulted in a tighter concentration of a single cluster. A similar tendency was observed when *h*^2^ = 0.2 supporting the findings that the extent of measures of connectedness between individuals from different management units is enhanced with genomic information. The remaining PC plots for PEVD (**A** and **G**) and r (**A** and **G_s_**) are in Figures S3, S4, S5, and S6 in File S1, which presented a pattern similar to Figures 4 and 5.

**Figure 4:**
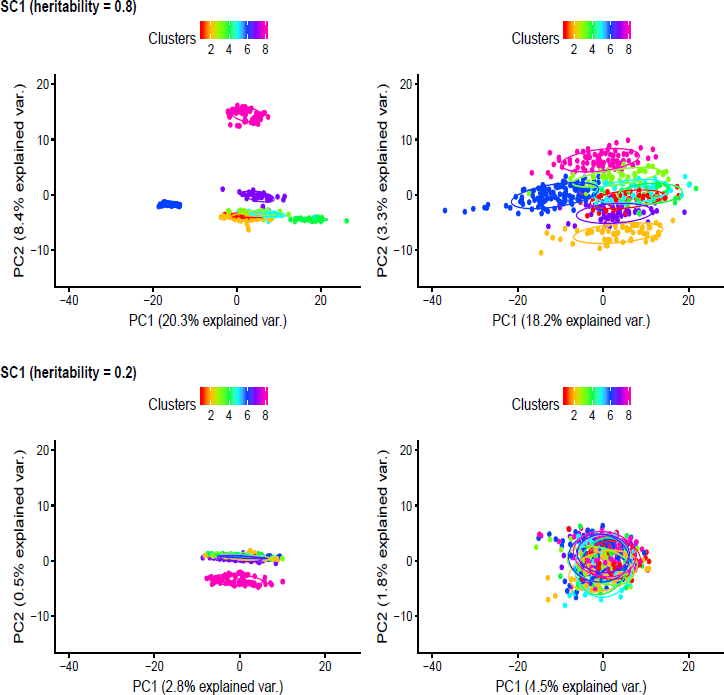
Principal component (PC) plots for Scenario 1 with coefficient of determination (CD) statistics. The first and second rows are according to heritability of 0.8 and of 0.2. The first and second columns are derived from pedigree-based (**A**) and genome-based (**G**) CD, respectively. The PC plots were grouped by clusters and colored in different colors. Individuals within the same cluster were grouped by the circles.

**Figure 5:**
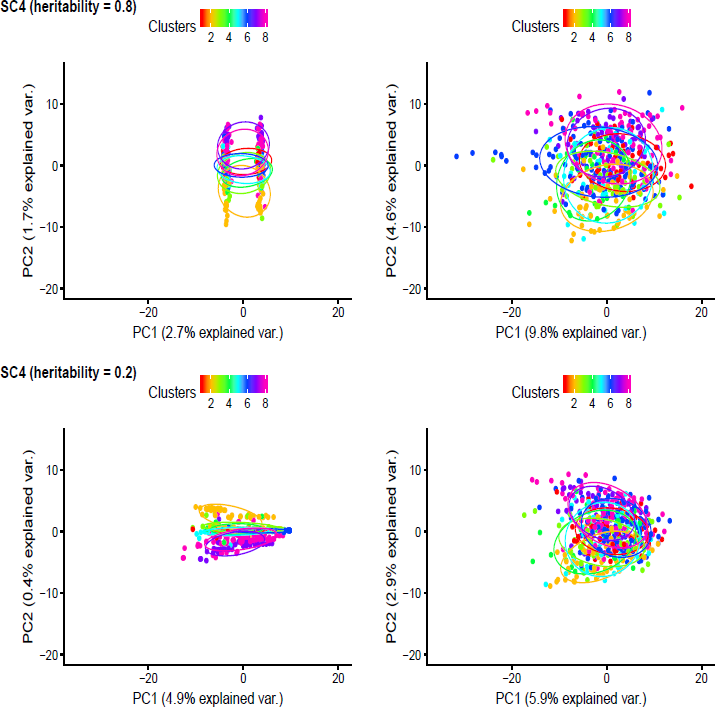
Principal component (PC) plots for Scenario 4 with coefficient of determination (CD) statistics. The first and second rows are according to heritability of 0.8 and of 0.2. The first and second columns are derived from pedigree-based (**A**) and genome-based (**G**) CD, respectively. The PC plots were grouped by clusters and colored in different colors.
Individuals within the same cluster were grouped by the circles.

## Discussion

With sufficient connectedness across management units, BLUP of genetic values can be fairly compared. Without such connectedness, making selection decisions based on breeding values of individuals from different management units might be associated with an increased risk of uncertainty in genetic evaluation due to imperfect separation of the genetic signal from noise. In addition to PEVD, CD, and r, other connectedness measures have been applied to pedigree data (e.g., Foulley et al., 1992; Fouilloux et al., 2008), which have their own characteristics. Advancement of molecular biotechnology now enables us to assess connectedness at the genomic level. Although genomic data are clearly important in genetic evaluations due to increased accuracy of estimates of genetic merit for non-parent individuals, little consideration has been given to the effect of genomic information on connectedness measures. In this study, we employed three measures of connectedness to examine the extent to which genomic information increases the estimates of connectedness.

## Relatedness in quantitative genetics

The majority of connectedness among management units was driven by the degree of genetic links or relatedness. The theory behind relatedness is largely entrenched in quantitative genetics dating back to work of Fisher (1918) and Wright (1921). Quantitative genetics offers a useful framework to study traits and diseases that are controlled by a considerable number of small effect genes. For traits with polygenic genetic architectures, inheritance does not exhibit a clear genotype-phenotype pattern. However, genetic resemblance between relatives (e.g., the genetic correlations between parent and offspring or between pairs of different types of siblings) can be exploited to estimate quantitative genetic parameters. For this reason, genetic resemblance between relatives has been at the heart of quantitative genetics. Consequently, the vast majority of the theoretical developments and applications of the last century were built around family data. The availability of dense panels of common SNPs has made it possible to trace Mendelian sampling and hence capture more detailed relatedness compared to pedigree information. It enables quantifying genomic kinships among related individuals that are not otherwise apparent because of incomplete pedigrees or the general assumption that animals in a baseline or founder population are unrelated. Thus, it has opened new opportunities for quantitative genetic analysis using data from distant relatives. The rationale is that individuals are genomically related to some extent and molecular similarity introduces covariance even if individuals are not related in the sense of known pedigree. These factors possibly contribute to the reduction of PEV or increase of PEC and hence lead to increased capturing of genetic connectedness in PEVD, CD, and r such that genetic merit estimates can be better compared across management units.

## The impact of genomic information on connectedness

We found from the mice data that genomic information increased favorable changes in measures of connectedness among individuals from different management units and reduced the risk of potential uncertainty in EBV-based comparisons when selecting individuals across management units. In addition, the rate of improvement in measures of connectedness in PEVD and r was greater when there was at least one full-sib in different management units. This is in concordance with Legarra et al. (2008) who used the same dataset and reported that the use of genome-wide selection increased predictive performance up to 0.22 across families and up to 0.03 within families compared to pedigree-based regression counterparts. On the other hand, CD accounted for the reduction of variability of relatedness between individuals under comparison resulting in decreased estimates of connectedness. Analysis of cattle data supported the results from mice and revealed that the benefit of using genomic information is greater for a disconnected design rather than a connected design. PCA was performed to visualize improvement in connectedness when moving from pedigree to genomic-based relationships. The PC plots supported the evidence that genomic information can improve detection of connectedness between individuals from different management units. This is particularly so when more individuals from the same clusters are assigned to different management units.

## Choice of kernel matrices

Unlike PEVD and CD, comparisons between the **A** and **G** kernel matrices evaluated by the r statistic behaved irregularly. By examining the specific components of the PEV matrix for **G** and **A** in the mice dataset, we found genomic information reduces off-diagonal elements more than diagonals. This illustrates a fundamental difference between r and either PEVD or CD because this statistic is based on the ratio, rather than the magnitude of individual elements. It may be argued that the inconsistent connectedness results from r occur because the **G** matrix is not on the same scale as the **A** matrix, suggesting that r statistics are not invariant regarding how genomic relatedness is defined. Given pedigree information, the numerator relationship matrix is defined as IBD. On the other hand, given a marker matrix, there are a number of ways to construct a genomic relationship matrix as discussed by Toro et al. (2011). The G matrix we used captures the proportion of the genome that is IBS by accounting for the covariance structure among individuals by molecular markers (Toro et al., 2002). This kernel matrix is an estimator of IBD relationships (Powell et al., 2010). Caution should be exercised when interpreting connectedness measures derived using genomic data as the underlying assumption is that relationships are built based on alleles being IBS and not necessarily being IBD. Therefore, we attempted to make **G** more compatible to **A** by using **G_0.5_** derived from allele frequencies equal to 0.5 and by using the min-max scaler transformation to produce the scaled genomic relationship matrix **G_s_**. For instance, compared to using **G**, entries of PEV matrix from using **G_s_** were more similar to those **A**, especially when there was connectedness, and in turn r statistics yielded greater connectedness values. Although connecting marker-based genomic relatedness to classical theory is still an open question in quantitative genetics, care needs to be taken when comparing genetic connectedness with genomic connectedness especially when the ratio-based statistic is used. Moreover, many additional factors may influence the elements of IBS matrix such as the choice of MAF, the density of SNPs, imperfect linkage disequilibrium (LD) between markers and quantitative trait loci, and error associated with estimating genomic relationships from a finite set of markers (e.g., Goddard, 2009).

## Choice of connectedness statistics

There was an issue with PEVD coupled with the **G_s_** matrix in the cattle dataset when *h*^2^ was 0.2 as the estimates of connectedness were less than those using **A** (Table 3). Note that this was not the case when *h*^2^ was equal to 0.8. The **H** matrix blended from the **A** and **G** kernel matrices yielded the estimates of connectedness that lie somewhere between the results obtained when using **A** and **G** alone. However, this pattern was not observed when **G_s_** was used in conjunction with **A** to compute PEVD (Table S1 in File S1). Apparently, scaling has a negative influence on blending for PEVD, which warrants further research. One potential reason with **G_s_** for the discrepancy is the proportional increase of PEC relative to PEV is larger when transitioning from the **A** to **G_s_**. This issue of proportional change is similar to that observed earlier with the r statistic coupled with **G**. These results illustrate that connectedness statistics are not invariant with respect to how the genomic relationship matrix is created and each of them captures different aspects of genomic connectedness. The CD was the only one statistic that yielded consistent estimates of increased connectedness throughout this study. Its consistency was observed regardless of choice of kernel matrices, heritability levels, datasets used, and simulated scenarios for management units.

**Table 3:** Average genetic connectedness statistics across management units in the cattle data. S1 (completely disconnected), S2 (disconnected), S3 (partially connected), and S4 (connected) represent four management unit scenarios. PEVD, CD, and r denote prediction error variance of the difference, coefficient of determination, and prediction error correlation. We compared pedigree-based **A**, standard genome-based **G**, genome-based **G_0.5_** assuming equal allele frequencies, and scaled genome-based **G_s_** kernel matrices to evaluate relationships among individuals. Two heritability values 0.8 and 0.2 were simulated.

| Scenarios | Methods | Kernels | Heritability ($h^2$) | |
| --- | --- | --- | --- | --- |
|  |  |  | 0.8 | 0.2 |
| S1 | PEVD | $\mathbf{A}$ | 0.077 | 0.102 |
| | | $\mathbf{G}$ | 0.051 | 0.085 |
| | | $\mathbf{G}_{0.5} (\mathbf{G}_s)$ | 0.039 (0.066) | 0.066 (0.110) |
| | CD | $\mathbf{A}$ | 0.324 | 0.112 |
| | | $\mathbf{G}$ | 0.539 | 0.224 |
| | | $\mathbf{G}_{0.5} (\mathbf{G}_s)$ | 0.528 (0.558) | 0.195 (0.265) |
| | $r_{ij}$ | $\mathbf{A}$ | 0.017 | 0.005 |
| | | $\mathbf{G}$ | -0.014 | -0.007 |
| | | $\mathbf{G}_{0.5} (\mathbf{G}_s)$ | 0.725 (0.468) | 0.465 (0.174) |
| S2 | PEVD | $\mathbf{A}$ | 0.016 | 0.022 |
| | | $\mathbf{G}$ | 0.011 | 0.020 |
| | | $\mathbf{G}_{0.5} (\mathbf{G}_s)$ | 0.008 (0.014) | 0.015 (0.025) |
| | CD | $\mathbf{A}$ | 0.376 | 0.152 |
| | | $\mathbf{G}$ | 0.636 | 0.326 |
| | | $\mathbf{G}_{0.5} (\mathbf{G}_s)$ | 0.625 (0.652) | 0.290 (0.373) |
| | $r_{ij}$ | $\mathbf{A}$ | 0.014 | 0.004 |
| | | $\mathbf{G}$ | -0.015 | -0.007 |
| | | $\mathbf{G}_{0.5} (\mathbf{G}_s)$ | 0.738 (0.496) | 0.468 (0.177) |
| S3 | PEVD | $\mathbf{A}$ | 0.012 | 0.018 |
| | | $\mathbf{G}$ | 0.008 | 0.016 |
| | | $\mathbf{G}_{0.5} (\mathbf{G}_s)$ | 0.007 (0.011) | 0.013 (0.020) |
| | CD | $\mathbf{A}$ | 0.460 | 0.211 |
| | | $\mathbf{G}$ | 0.653 | 0.346 |
| | | $\mathbf{G}_{0.5} (\mathbf{G}_s)$ | 0.649 (0.669) | 0.312 (0.394) |
| | $r_{ij}$ | $\mathbf{A}$ | 0.018 | 0.005 |
| | | $\mathbf{G}$ | -0.012 | -0.006 |
| | | $\mathbf{G}_{0.5} (\mathbf{G}_s)$ | 0.739 (0.498) | 0.468 (0.178) |
| S4 | PEVD | $\mathbf{A}$ | 0.007 | 0.007 |
| | | $\mathbf{G}$ | 0.005 | 0.007 |
| | | $\mathbf{G}_{0.5} (\mathbf{G}_s)$ | 0.004 (0.007) | 0.005 (0.009) |
| | CD | $\mathbf{A}$ | 0.125 | 0.048 |
| | | $\mathbf{G}$ | 0.367 | 0.132 |
| | | $\mathbf{G}_{0.5} (\mathbf{G}_s)$ | 0.362 (0.384) | 0.114 (0.158) |
| | $r_{ij}$ | $\mathbf{A}$ | 0.024 | 0.008 |
| | | $\mathbf{G}$ | -0.007 | -0.002 |
| | | $\mathbf{G}_{0.5} (\mathbf{G}_s)$ | 0.741 (0.502) | 0.470 (0.181) |

## Future direction

One important direction for future study is to investigate whether increased connectedness observed by genomic relatedness also leads to increased predictive accuracy of genetic values across management units assessed by cross-validation. In this case, across management units can be considered as training and testing sets. In addition, while the current norm of genomic prediction is to use an IBS relationship matrix that aims to capture relationships at unknown QTLs through LD between markers and QTLs, we argue that improving the quality of breeding value comparisons and improving the accuracy of genomic prediction can be viewed as relevant but two different items. In this regard, a genome-wide IBD relationship matrix (e.g., Fernando and Grossman, 1989), where marker inheritance is traced through a known pedigree, may be worthwhile to revisit for the purpose of ascertaining connectedness in a future study.

Also, for the r statistic, we summarized connectedness by averaging the r statistic of pairs of individuals across units rather than by averaging the relevant components of PEC and PEV followed by taking their ratio; our justification for that choice was provided in the Appendix. When the latter summary statistic was used for r, the differences were negligible in the mice data and the pattern was the same for the scenario 1 of cattle data.

In conclusion, this study confirms that use of genomic relatedness improved genetic connectedness across management units compared to use of pedigree relationships. To our knowledge, this marks the first thorough investigation of genomic connectedness. We contend that our work is a critical first step toward better understanding genetic connectedness that may have a positive impact on genomic evaluation of agricultural species.

## Acknowledgments

This work was supported in part by the University of Nebraska startup funds to G.M. The authors thank Dale Van Vleck and Larry Kuehn for their valuable comments on the manuscript.

## Appendix

### Prediction error correlation statistic across units

When a population is divided into two management units, and relatedness between those two units is based on the **G** matrix, the flock connectedness or unit connectedness correlation (r) of Kuehn et al. (2008) always yields an estimate of -1. Here, we wish to illustrate that result. Flock connectedness is derived by averaging the relevant components of PEC and PEV followed by taking their ratio. Suppose that there are two units and the numbers of individuals in units *i′* and *j′* are *n_i′_* and *n_j′_*, respectively. The total number of individuals is *N* = *n_i′_* + *n_j′_*. The assumptions of the G matrix is that the genotyped individuals represent the base population where the expected value of self-relatedness is 1, assuming no inbreeding, and that the mean relatedness of any one individual to the rest of the individuals is zero. Consequently, the expectation of diagonal elements of the G matrix is equal to the number of individuals assuming no inbreeding and the expectation of off-diagonal elements is –1/(*N* – 1) (e.g., Yang et al., 2013). Then the expectation of numerator in the r statistic is proportional to

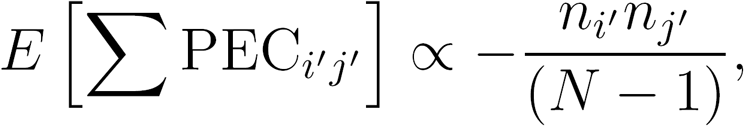

and the expectation of denominator is the square root of the product between

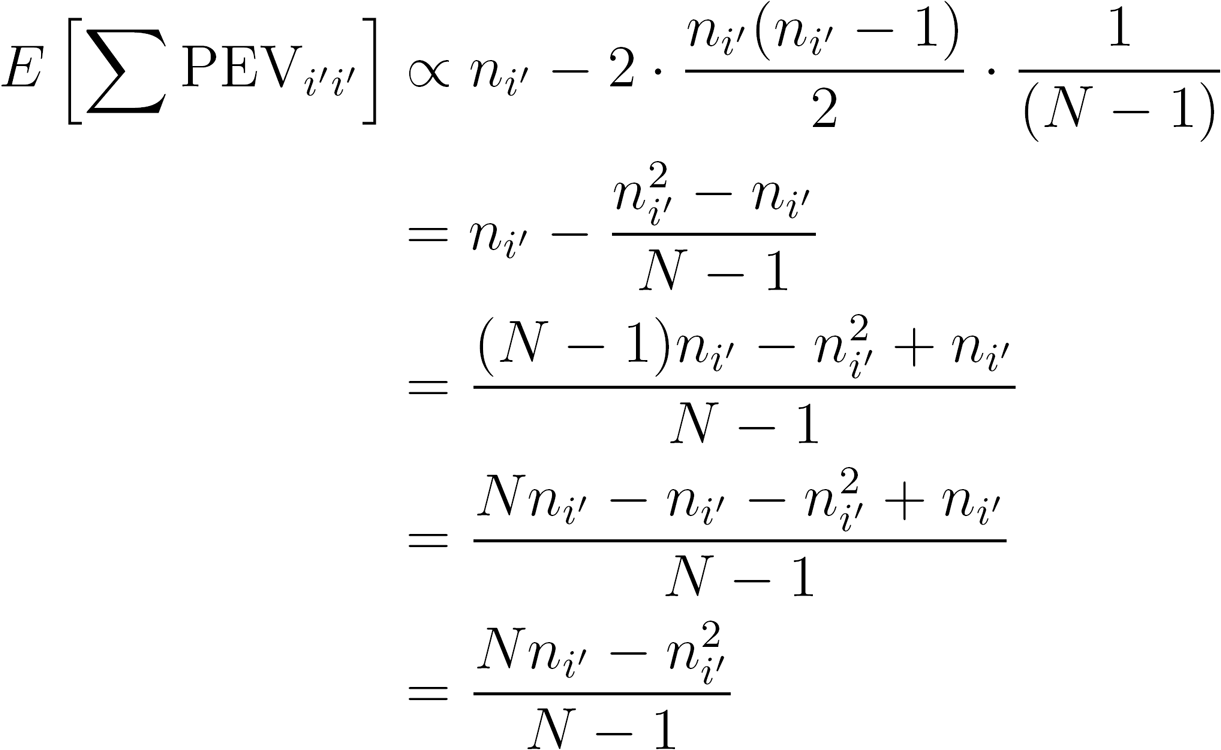

and

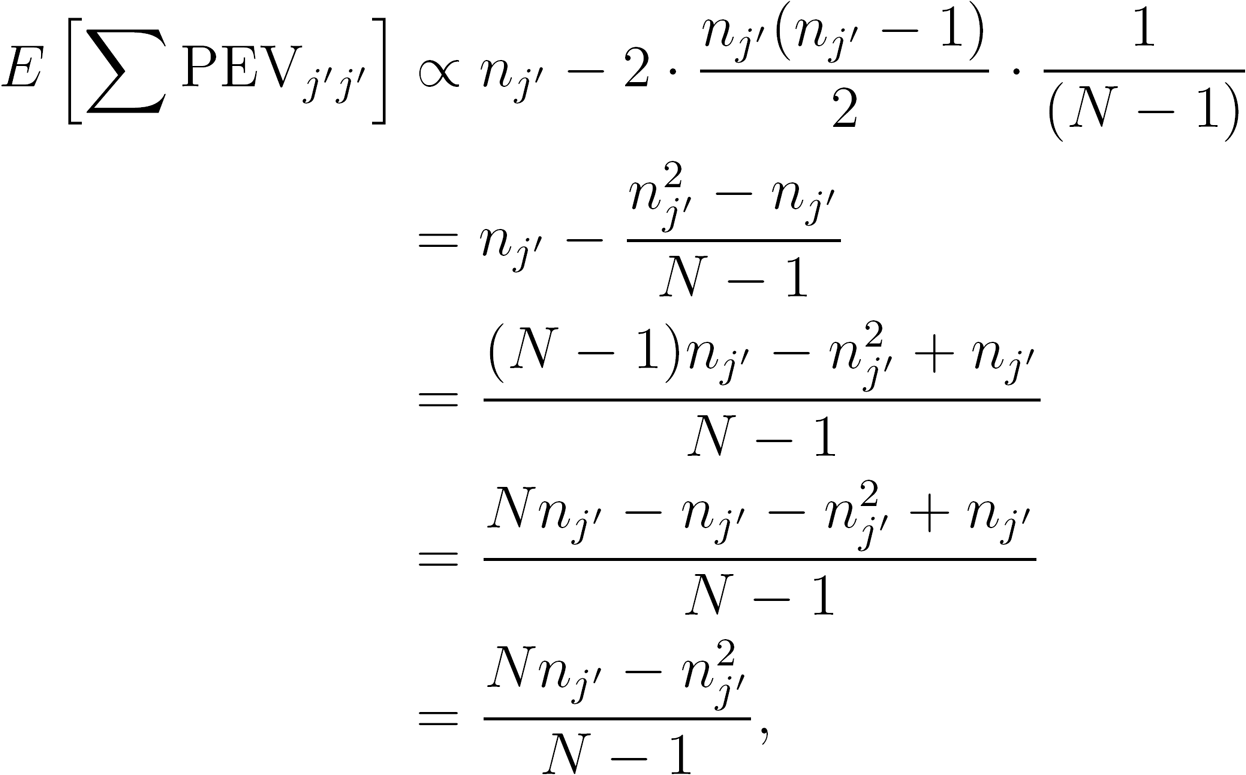

so that

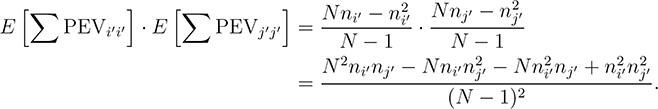

Note that the first three terms in the numerator are equal to zero

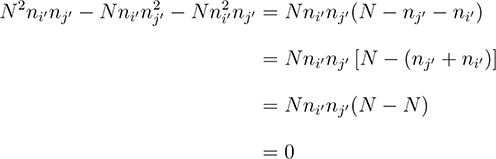

because *N* = *n_i′_*+*n_j′_*. Therefore, the r statistic between units *i*′ and *j*′ is given by

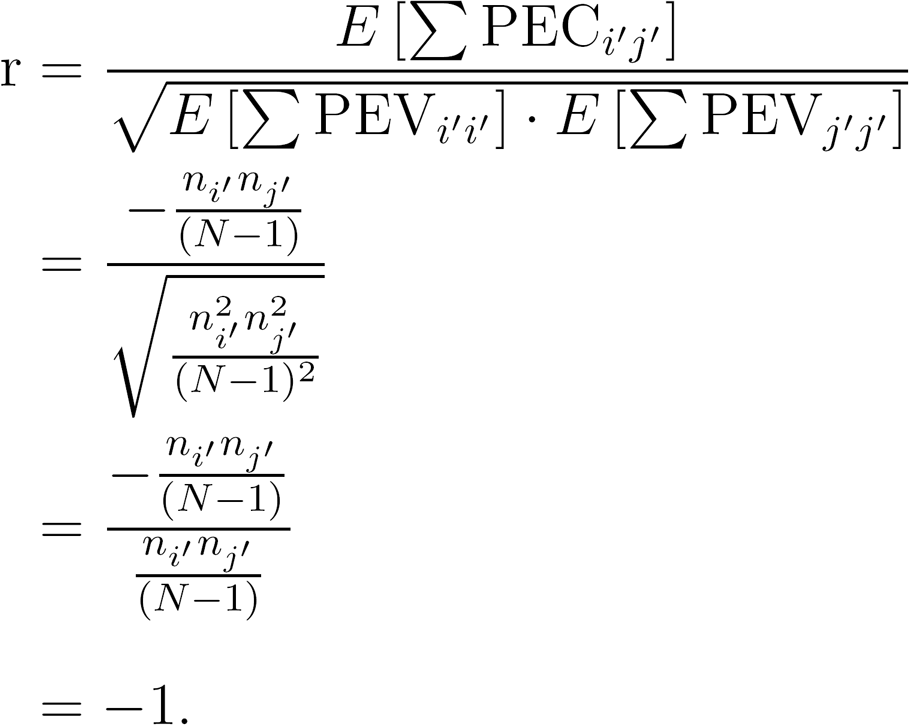

When *N* = *n_i′_* + *n_j′_*, this result holds regardless of relatedness level, connectedness level, and how individuals are partitioned into the two management units *i′* and *j′*. The partitioning of animals into two distinct units is particularly relevant in the context of genomic prediction where animals may be divided into training and testing sets. In this scenario, computing the connectedness between the two sets along the lines of Rincent et al. (2012) and Isidro et al. (2015) is potentially informative relative to expectations of the performance of resulting genomic predictors. The **G_s_** or **G_0.5_** matrix changes the expectation of off-diagonal elements to positive values and shifts the statistic by a constant as explained in the Methods section, yielding connectedness between units *i′* and *j′* of close to 1. Because scenarios 2 to 4 in the cattle dataset simulated two management units, the average of the r-statistic of pairs of individuals in different management units was used to summarize connectedness in this study. Note that this is shown as lamb connectedness or individual connectedness in Kuehn et al. (2008). The two types of connectedness differ mainly by whether we take the average followed by the ratio (unit connectedness) or take the ratio first followed by the average (individual connectedness).

## References

Christensen, O. F. and Lund, M. S. (2010). Genomic prediction when some animals are not genotyped. Genet Sel Evol., 42:2.

Eikje, L.S. and Lewis, R. M. (2015). Strong connectedness within Norwegian Cheviot and Fur Sheep ram circles allows reliable estimation of breeding values. J Anim Sci., 93:3322–3330.

Fernando, R. and Grossman, M. (1989). Marker assisted selection using best linear unbiased prediction. Genet Sel Evol., 21:467–477.

Fisher, R.A. (1918). The correlation between relatives on the supposition of Mendelian inheritance. Trans. R. Soc. Edin, 52:399–433.

Fouilloux, M.N., Clément, V., and Laloë, D. (2008). Measuring connectedness among herds in mixed linear models: from theory to practice in large-sized genetic evaluations. Genet Sel Evol., 40:145–159.

Foulley, J.L., Hanocq, E., and Boichard, D. (1992). A criterion for measuring the degree of connectedness in linear models of genetic evaluation. Genet Sel Evol., 24:315–330.

Goddard, M. (2009). Genomic selection: prediction of accuracy and maximisation of long term response. Genetica, 136:245–257.

Henderson, C.R. (1984). Applications of Linear Models in Animal Breeding. University of Guelph;, Third edition, Edited by Schaeffer LR. Guelph.

Hill, W.G. and Weir, B. S. (2011). Variation in actual relationship as a consequence of mendelian sampling and linkage. Genetics Research, 93:47–64.

Isidro, J., Jannink, J.-L., Akdemir, D., Poland, J., Heslot, N., and Sorrells, M. E. (2015). Training set optimization under population structure in genomic selection. Theor Appl Genet, 128:145.

Kaufman, L. and Rousseeuw, P. J. (1990). Finding groups in data: an introduction to cluster analysis. John Wiley and Sons, New York.

Kennedy, B.W. and Trus, D. (1993). Considerations on genetic connectedness between management units under an animal model. J Anim Sci., 71:2341–2352.

Kuehn, L.A., Lewis, R. M., and Notter, D. R. (2007). Managing the risk of comparing estimated breeding values across flocks or herds through connectedness: a review and application. Genet Sel Evol., 39:225.

Kuehn, L.A., Lewis, R. M., and Notter, D. R. (2009). Connectedness in Targhee and Suffolk flocks participating in the United States national sheep improvement program. J Anim Sci., 87:507–515.

Kuehn, L.A., Notter, D. R., Nieuwhof, G. J., and Lewis, R. M. (2008). Changes in connectedness over time in alternative sheep sire referencing schemes. J Anim Sci., 86:536–544.

Laloë, D. (1993). Precision and information in linear models of genetic evaluation. Genet Sel Evol., 25:557.

Laloë, D., Phocas, F., and Ménissier, F. (1996). Considerations on measures of precision and connectedness in mixed linear models of genetic evaluation. Genet Sel Evol., 28:359.

Legarra, A., Aguilar, I., and Misztal, I. (2009). A relationship matrix including full pedigree and genomic information. J. Dairy Sci., 92:4656–4663.

Legarra, A., Robert-Granié, C., Manfredi, E., and Elsen, J.-M. (2008). Performance of genomic selection in mice. Genetics, 180:611–618.

Lewis, R.M., Crump, R. E., Simm, G., and Thompson, R. (1999). Assessing connectedness in across-flock genetic evaluations. In Proc. Br. Soc. Anim. Sci., page 121, Scarborough, UK.

Maenhout, S., De Baets, B., and Haesaert, G. (2010). Graph-based data selection for the construction of genomic prediction models. Genetics, 185:1463–1475.

Momen, M., Mehrgardi, A. A., Sheikhy, A., Esmailizadeh, A., Fozi, M. A., Kranis, A., Valente, B. D., Rosa, G. J., and Gianola, D. (2017). A predictive assessment of genetic correlations between traits in chickens using markers. Genet Sel Evol., 49:16.

Powell, J.E., Visscher, P. M., and Goddard, M. E. (2010). Reconciling the analysis of ibd and ibs in complex trait studies. Nat Rev Genet., 11:800–805.

Pszczola, M., Strabel, T., van Arendonk, J. A. M., and Calus, M. P. L. (2012). The impact of genotyping different groups of animals on accuracy when moving from traditional to genomic selection. J Dairy Sci., 95:5412–5421.

Reynolds, A.P., Richards, G., de la Iglesia, B., and Rayward-Smith, V. J. (2006). Clustering rules: A comparison of partitioning and hierarchical clustering algorithms. J Math Model Algor., 5:475.

Rincent, R., Laloë, D., Nicolas, S., Altmann, T., Brunel, D., Revilla, P., Rodríguez, V. M., Moreno-Gonzalez, J., Melchinger, A., Bauer, E., Schoen, C. C., Meyer, N., Giauffret, C., Bauland, C., Jamin, P., Laborde, J., Monod, H., Flament, P., Charcosset, A., and Moreau, L. (2012). Maximizing the reliability of genomic selection by optimizing the calibration set of reference individuals: comparison of methods in two diverse groups of maize inbreds (Zea mays L.). Genetics, 192:715–728.

Solberg, L.C., Valdar, W., Gauguier, D., Nunez, G., Taylor, A., Burnett, S., Arboledas-Hita, C., Hernandez-Pliego, P., Davidson, S., Burns, P., Bhattacharya, S., Hough, T., Higgs, D., Klenerman, P., Cookson, W. O., Zhang, Y., Deacon, R. M., Rawlins, J. N. P., Mott, R., and Flint, J. (2006). A protocol for high-throughput phenotyping, suitable for quantitative trait analysis in mice. Mamm Genome, 17:129–146.

Toro, M., Barragán, C., Óvilo, C., Rodrigañez, J., Rodriguez, C., and Siliá, L. (2002). Estimation of coancestry in Iberian pigs using molecular markers. Conserv Genet., 3:309–320.

Toro, M.A., García-Cortés, L. A., and Legarra, A. (2011). A note on the rationale for estimating genealogical coancestry from molecular markers. Genet Sel Evol., 43:27.

Valdar, W., Solberg, L. C., Gauguier, D., Burnett, S., Klenerman, P., Cookson, W. O., Taylor, M. S., Rawlins, J. N. P., Mott, R., and Flint, J. (2006). Genome-wide genetic association of complex traits in heterogeneous stock mice. Nat Genet, 38:879–887.

VanRaden, P.M. (2007). Genomic measures of relationship and inbreeding. Interbull Bull., 37:3336.

VanRaden, P.M. (2008). Efficient methods to compute genomic predictions. J. Dairy Sci., 91:4414–4423.

Vitezica, Z.G., Aguilar, I., Misztal, I., and Legarra, A. (2011). Bias in genomic predictions for populations under selection. Genet Res., 93:357–366.

Wimmer, V., Albrecht, T., Auinger, H.-J., with contributions by Malena Erbe, C.-C. S., Ober, U., and Reimer, C. (2015). synbreedData: Data for the Synbreed Package. R package version 1.5.

Wright, S. (1921). Systems of mating. I. The biometric relations between offspring and parent. Genetics, 6:111–123.

Wright, S. (1922). Coefficients of inbreeding and relationship. Am Nat., 56:330–338.

Yang, J., Lee, S. H., Goddard, M. E., and Visscher, P. M. (2013). Genome-wide complex trait analysis (gcta): methods, data analyses, and interpretations. Genome-wide association studies and genomic prediction, pages 215–236.

